# Does Vaping Increase the Likelihood of SARS-CoV-2 Infection? Paradoxically Yes and No

**DOI:** 10.1101/2022.09.09.507373

**Authors:** Rattapol Phandthong, Man Wong, Ann Song, Teresa Martinez, Prue Talbot

**Affiliations:** Department of Molecular, Cell and System Biology, University of California, Riverside, CA 92521, USA

**Keywords:** e-cigs, SARS-CoV-2, COVID-19, ACE2, TMPRSS2, Air-liquid-Interface

## Abstract

Data on the relationship between electronic cigarettes (ECs) and SARS-CoV-2 infection are limited and contradictory. Evidence indicates that EC aerosols or nicotine increase ACE2, SARS-CoV-2 virus receptors, which increase virus binding and susceptibility. Our objectives were to determine if EC aerosols increased SARS-CoV-2 infection of human bronchial epithelial cells and to identify the causative chemical(s). A 3D organotypic model (EpiAirway™) in conjunction with air liquid interface (ALI) exposure was used to test the effects of aerosols produced from JUUL™ “Virginia Tobacco” and BLU™ ECs, or individual chemicals (nicotine, propylene glycol, vegetable glycerin (PG/VG), and benzoic acid) on infection using SARS-CoV-2 pseudoparticles. Exposure of EpiAirway™ to JUUL™ aerosols increased ACE2, while BLU™ and lab-made EC aerosols containing nicotine increased ACE2 levels and TMPRSS2 activity, a spike protease that enables viral-cell fusion. Pseudoparticle infection of EpiAirway™ increased with aerosols produced from PG/VG, PG/VG plus nicotine, or BLU™ ECs. JUUL™ EC aerosols did not increase infection above controls. The baseline level of infection in JUUL™ treated aerosol groups was attributed to benzoic acid, which mitigated the enhanced infection caused by PG/VG or nicotine. The benzoic acid protection from enhanced infection continued at least 48 hours after exposure. TMPRSS2 activity was significantly correlated with e-liquid pH, which in turn was significantly correlated with infection, with lower pH blocking PG/VG and nicotine-induced-enhanced infection. While ACE2 levels increased in EpiAirway™ tissues exposed to EC aerosols, infection depended on the ingredients of the e-liquids. PG/VG and nicotine enhanced infection, an effect that was mitigated by benzoic acid.

## Introduction

Electronic cigarettes (ECs) are nicotine delivery devices that aerosolize e-liquids, which usually contain nicotine, propylene glycol (PG), vegetable glycerin (VG), and flavor chemicals [1,2]. While sometimes promoted as less harmful than tobacco cigarettes, ECs are not harm free [3-7]. Adverse effects reported for EC users include impaired pulmonary immune defenses against bacterial and viral infections [8] and a weakened innate immune response in the lung compared to non-users [9]. Despite their popularity, especially among adolescents [10], little work has been done on the relationship between EC use and users’ susceptibility to infection by SARS-CoV-2, which causes Coronavirus Disease 2019 (COVID-19). The entry of SARS-CoV-2 into host cells depends on the viral spike protein receptor (angiotensin converting enzyme-2 or ACE2) and the cellular transmembrane protease, serine 2 (TMPRSS2), which cleaves an internal site of the spike S2 subunit preparatory to fusion [11, 12].

A survey of adolescents reported that EC users were five times more likely to be diagnosed with COVID-19 than non-users [13], and a correlative study found a positive association between EC use and SARS-CoV-2 cases [14]. Whole body exposure of mice to EC aerosol with or without nicotine elevated ACE2 in the lung [15-18], suggesting an increased in the likelihood of infection. Although human data are limited, Ghosh et al., 2022 [19] showed that EC aerosol increased ACE2 activity and soluble ACE2 levels in EC users’ bronchoalveolar lavage fluid; these changes were correlated with infection of human bronchial epithelial cells by SARS-CoV-2 pseudoparticles following exposure to JUUL™ “Virginia Tobacco” e-liquid in submerged cultures. Additionally, e-liquid with and without nicotine increased ACE2 expression in BEAS-2B cells after submerged treatment [20], an effect subsequently linked to nicotine concentration in air-liquid interface (ALI) exposures of BEAS-2B cells [21]. Using BEAS-2B monolayer cultures, we further showed that submerged and ALI exposures with nicotine and e-liquid with nicotine increased ACE2 levels, TMPRSS2 activity, and SARS-CoV-2 pseudoparticle infection [21]. While preliminary, these studies suggest that EC use may increase the likelihood of contracting COVID-19. In contrast, hospital observational studies have concluded that EC use is not associated with an increased risk for COVID-19 [23, 24].

Because data on EC use and SARS-CoV-2 susceptibility are limited and contradictory, our objective was to test the hypothesis that using ECs is a risk factor for SAR-CoV-2 infection using controlled laboratory experiments. The current study is the first to examine SARS-CoV-2 infection using 3D organotypic cultures of human bronchial epithelium (EpiAirway™) exposed at the ALI. EpiAirway™ tissues, which contain ciliated, basal, and mucus producing cells, are the best in vitro model available for examining how EC use affects SAR-CoV-2 infection. To elucidate the effect of vaping on COVID-19 and to identify the specific chemicals in EC liquids that affect SARS-CoV-2 infection, authentic JUUL™ or BLU™ EC aerosols and aerosols made with specific e-liquid ingredients (nicotine and PG/VG or mixtures thereof) were tested using ALI exposures of EpiAirway™ tissues. Infection was evaluated using SARS-CoV-2 viral pseudoparticles containing spike protein and a green-fluorescent reporter. The three endpoints examined for each exposure treatment included: (1) ACE2 levels in EpiAirway™ tissues; (2) TMPRSS2 protease activity; and (3) SARS-CoV-2 viral pseudoparticle infection. Our data show that the relationship between vaping and SARS-CoV-2 infection is complex and highly dependent on the ingredients of the e-liquid used to create the aerosol. These data will provide EC users with options that may reduce their risk of contracting COVID-19, affect future research studies, be of interest to regulatory agencies, and help improve the design of clinical trials involving the use of tobacco products and SARS-CoV-2 infection.

## Materials and Methods

### Aerosol exposures at the ALI in the VITROCELL® cloud chamber

In brief, 3D EpiAirway™ tissues cultured on 12 well Transwell® inserts were placed in a VITROCELL® cloud chamber (VITROCELL®, Walkirch, Germany) for an ALI exposure to various chemical aerosols that were generated without heating. EpiAirway™ tissues were exposed to 5 puffs of PBS-, nicotine (0.3 mg/mL), or benzoic acid (0.2 mg/mL) aerosol. After the exposure, the cells were returned to the incubator to recover for 24 hrs. Deposition of nicotine was previously reported [24]. Additional details of cloud chamber exposure are in the supplement

### EC aerosol exposure at the ALI in the Cultex® RFS compact exposure system

A Cultex® RFS compact exposure module (Cultex Laboratories GmbH, Hannover, Germany) was used to expose cells or tissue cultures to humified sterile air (clean air control) or aerosols generated from EC devices. Prior to each exposure, cells or tissues samples were placed into the exposure chamber, which contained culture medium and was maintained at a temperature of 37 ºC. EpiAirway™ 3D tissues were exposed to 50 puffs of air or EC aerosols/day for 3 days. Between each exposure day, tissues were returned to the incubator and after the last exposure, tissues were allowed to recover in the incubator for 24 hrs prior to analyses. Deposition of nicotine was previously reported [24]. Additional details on Cultex® system exposures are in the Supplement.

### Quantification of TMPRSS2 proteolytic activity in EpiAirway™ tissues

The TMPRSS2 enzyme assay was performed using a previously published protocol, with some modification [24]. The fluorogenic substrate, Boc-Gln-Ala-Arg-AMC·HCl (Bachem, Torrance, CA. USA) was dissolved in DMSO and diluted in reaction buffer (50 mM Tris pH 8, 150 mM NaCl) to a final concentration of 10 μM. The fluorogenic substrate was added to each well of a 96-well plate, and fluorescence intensity was measured at 340/440 nm over 1 hr at 37 °C using a BioTek Synergy HTX (Agilent, Santa Clara, CA, USA).

### Spike viral pseudoparticle production

Pseudoparticle production was performed as outlined in Crawford et al., 2020 [25]. In brief, HEK293T cells were plated with antibiotic-free medium at a density of 7 × 10^6/^/T75 flask and transfected with Lipofectamine3000 (Thermo Fisher Sci, Waltham, MA, USA) using a total of 15 μg of lentiviral plasmids (described in the Supplement). Fluorescence microscopy was used to visually inspect transfection efficiency and the expression of the ZsGreen. The lentivirus was collected by centrifugation and the pellet was suspended in Viral Re-suspension Solution (Abcam, Cambridge, UK). Additional details of spike viral pseudoparticle are in the Supplement.

### Viral pseudoparticle infection

For each viral pseudoparticle infection experiment, a 0.3 multiplicity of infection (MOI) was used to infect 3D EpiAirway™ tissues. Viral pseudoparticles were delivered as a mixture with the appropriate fresh culture medium. After the recovery period, 100 μL of pseudoparticle medium was added directly onto the apical side of the Transwell®. Viral pseudoparticles were incubated with tissues for 24 hours, then the pseudoparticle mixture was removed. Tissues were allowed to incubate another 24 hours to amplify the expression of the green-fluorescent reporter protein in the infected cells. Cells were harvested and analyzed with flow cytometry to determine the number of infected cells. The flow cytometry protocol is in the Supplement.

**Additional Materials and Methods are available in the Supplement.**

## Results

### EC aerosol exposures at the ALI in the Cultex® system increased ACE2 in 3D EpiAirway™ tissues

The effects of authentic JUUL™ “Virginia Tobacco”, PG/VG, and PG/VG + nicotine aerosols were evaluated using 3D MatTek EpiAirway™ tissues, containing ciliated, mucus-producing, and basal cells (**Figure 1A)**. A 3-day EC aerosol exposure protocol was used to replicate an acute exposure that an EC user could receive. EpiAirway™ tissue was exposed to 50 puffs/day to be within the range (30-200 puffs day) reported for EC users [26].

**Figure 1.**
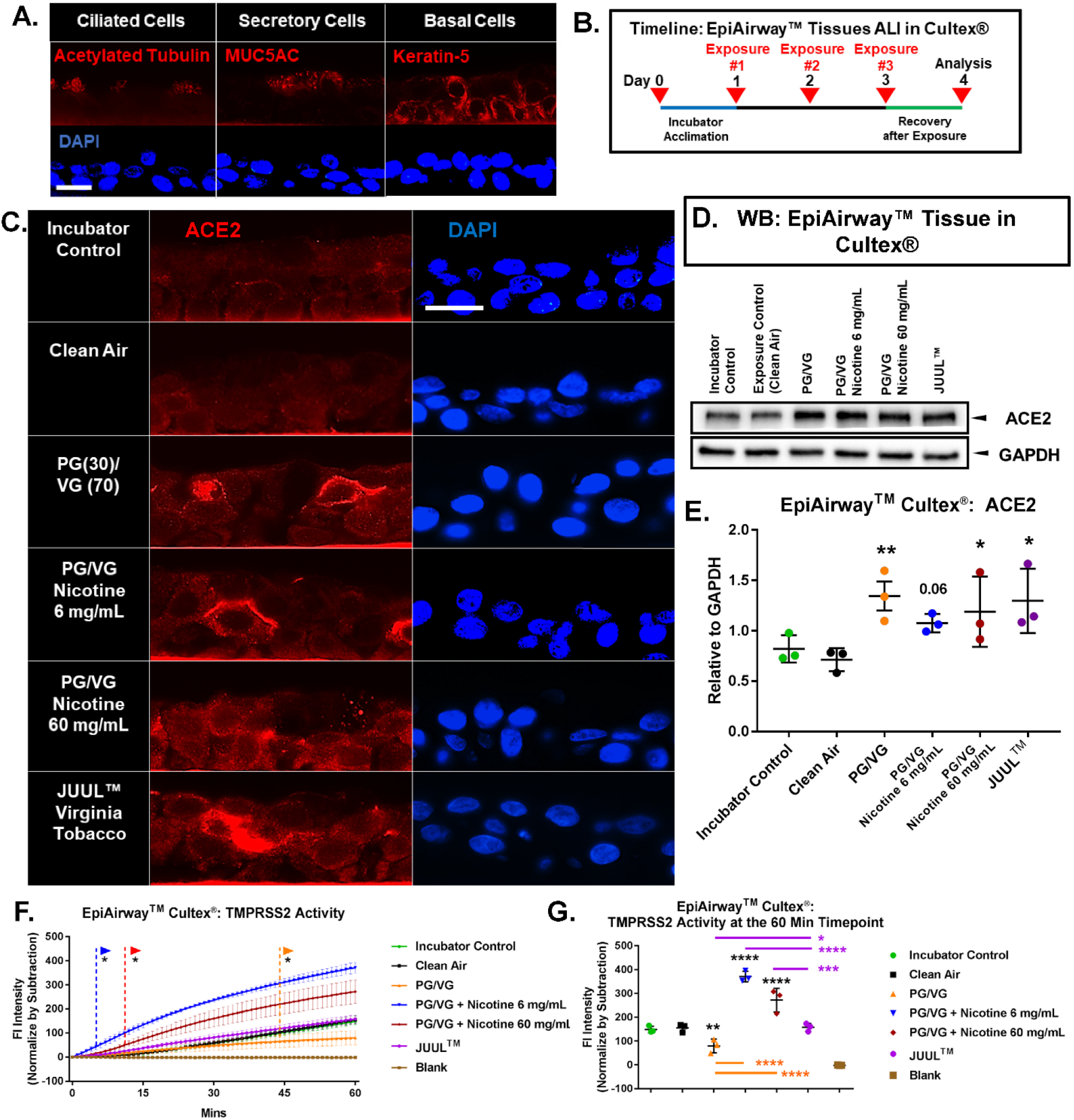
PG/VG, Nicotine, and JUUL™ EC aerosols affect ACE2 and TMPRSS2 in EpiAirway™ tissue following ALI exposure in a Cultex® system. **(A)** Micrographs showing cell types in EpiAirway™ tissues. Ciliated cells are labeled with acetylated tubulin. Secretory cells are labeled with MUC5AC. Basal cells are keratin-5 positive. Cell nuclei are shown by DAPI (blue) labeling. Scale bar = 50 μm **(B)** Experimental treatment used with cells exposed at the ALI in the Cultex® system. **(C)** Immunofluorescence labeling of EpiAirway™ tissues exposed to 50 puffs/day of EC aerosols for 3 days. Micrographs showing the tissue sections labeled with ACE2 antibody (red) and DAPI (blue). Scale bar = 50 μm **(D - E)** The effect of short-term EC aerosol exposures on ACE2 levels following ALI exposure in a Cultex® system (western blot). **(D)** A representative Western blot. (E) ACE2 levels relative to GAPDH averaged from three Western blots. **(F - G)** TMPRSS2 activity depends on EC aerosol formulations. (**F**) TMPRSS2 activity over 60 minutes. * indicate when values became significantly different than the control (*p* ranged from 0.05 to 0.0001). (**F**) was analyzed using a two-way ANOVA followed by Dunnett’s posthoc test to compare means to the exposure control. (**G**) Activity at the 60-minute timepoint for each treatment group. All graphs show the mean ± standard deviation of three independent experiments. In **E** and **G** Box-cox transformed data were analyzed using a one-way ANOVA followed by Tukey’s posthoc test. Black * = significantly different than the control. Color * = significantly different groups. *p < 0.05, **p < 0.01, ***p < 0.001, ****p < 0.0001.

The timeline in Figure 1B shows the experimental design and the sequence of EC aerosol exposure over 3 days. The EC aerosols were produced using authentic JUUL™ “Virginia Tobacco” pods or refillable pods containing either PG/VG only, PG/VG with 6 mg/mL of nicotine, or PG/VG with 60 mg/mL of nicotine. After the last exposure, tissues were returned to the incubator for 24 hours. They were then fixed for immunofluorescence microscopy **(Figure 1C)**, lysed for Western blot analysis **(Figure 1D-1E)**, or assayed for TMPRSS2 activity (**Figure 1 F-G**).

ACE2 receptors increased in all exposure groups except the clean air and incubator controls, which were similar to each other, showing that clean air exposure in the Cultex® system did not affect ACE2 levels **(Figure 1C)**. In Western blots, controls were not significantly different, but ACE2 levels increased significantly in all treatment groups compared to the clean air control **(Figure 1D-4E)**. These results indicate that acute exposure of EpiAirway™ to authentic JUUL™ aerosols, PG/VG, or PG/VG + nicotine can increase ACE2 levels, the SARS-CoV-2 point of entry.

### ALI exposure of EpiAirway™ tissues to EC aerosols with nicotine altered proteolytic activity of TMPRSS2

The activity of the spike-cleaving protease, TMPRSS2, was analyzed using a specific fluorogenic substrate **(Figure 1F-G)**. TMPRSS2 activity in EpiAirway™ tissues varied among treatment groups. Tissues exposed to PG/VG aerosols showed a small, but significant, decrease in TMPRSS2 activity when compared to the clean air control. EC aerosols containing PG/VG plus 6 or 60 mg/mL of nicotine significantly increased TMPRSS2 activity. In contrast, TMPRSS2 activity in the JUUL™ exposed group was significantly lower than tissue exposed to PG/VG with nicotine, even though JUUL™ fluid contained PG/VG and 60 mg/mL of nicotine. These data show the importance of testing mixtures and authentic EC aerosols, as well as measuring the enzymatic activity of TMPRSS2.

### Nicotine and PG/VG increased infection of EpiAirway™ tissues by viral pseudoparticles

Because EC aerosols increased ACE2 expression in EpiAirway™ tissues **(Figure 1C-1E)**, the following experiments were done to test the hypothesis that EC aerosols increase viral pseudoparticle infection. **Figure E1** shows the experimental timeline for short-term 3-day exposures of tissues followed by infection with SARS-CoV-2 viral pseudoparticles for 24 hours. Prior to flow cytometry assays, mock-infected and virus-infected cells were harvested, and the expression of ZsGreen in infected cells was verified using microscopy **(Figure 2A)**.

**Figure 2.**
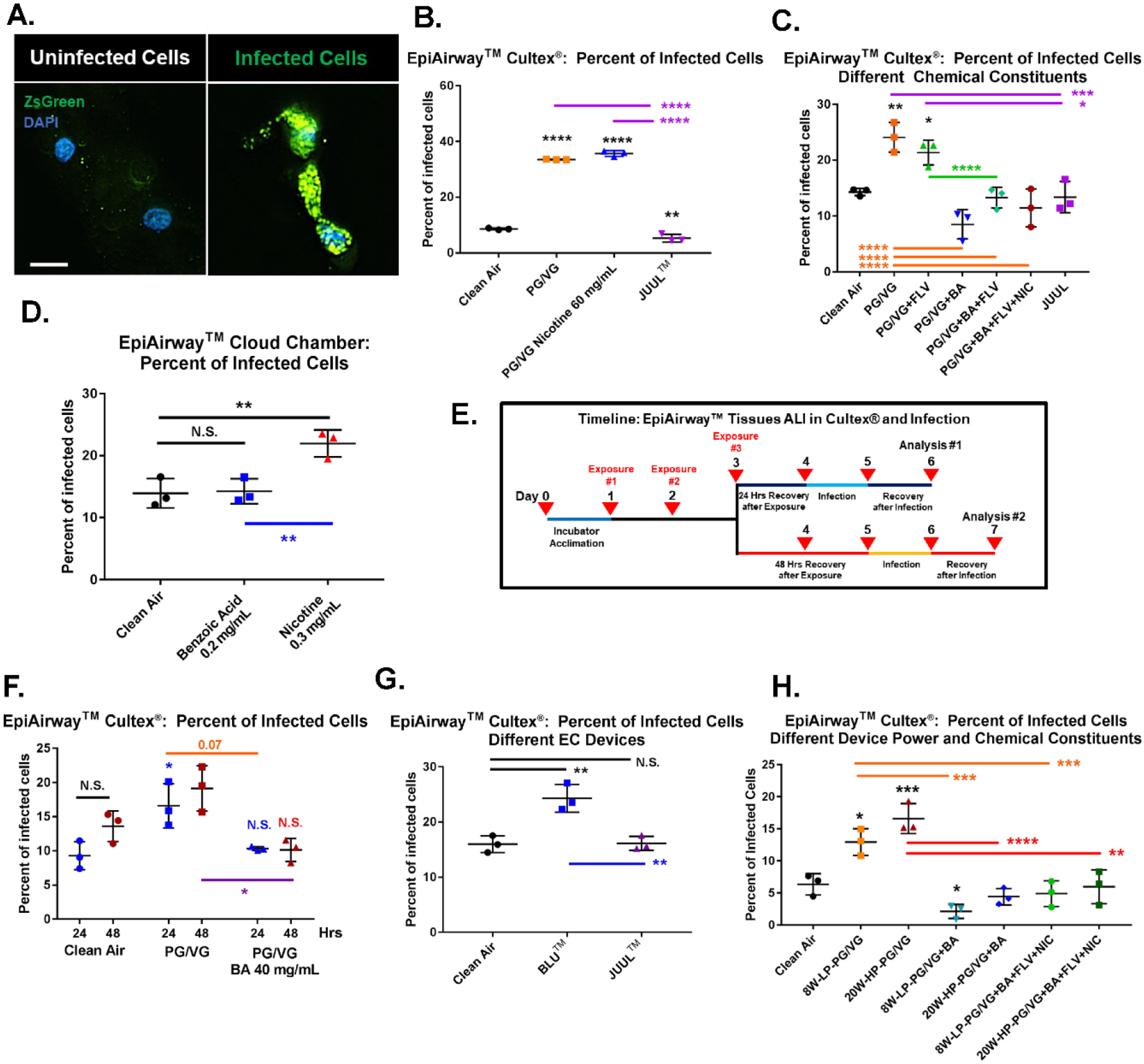
EC aerosols increased EpiAirway™ tissue susceptibility to viral pseudoparticle infection except when exposed to EC aerosols with benzoic acid. **(A)** Micrographs showing infected and uninfected cells with viral pseudoparticles. Infected cells expressed ZsGreen. Scale bar = 50 μm. **(B)** Flow cytometry data showing the percent of infected cells in the EpiAirway™ tissue after exposure to EC aerosols, followed by 24 hrs of exposure to pseudoparticles. PG/VG and PG/VG + nicotine 60 mg/mL increased infection, but JUUL™ “Virginia Tobacco” EC fluid did not. **(C)** Percent of infected cells after exposure to aerosols produced from different e-liquid formulations. Exposures were done in a Cultex® system using JUUL™ compatible pods filled with different mixtures. Infection did not increase in treatments containing benzoic acid. **(D)** Percent of infected cells after exposure to pure benzoic acid or nicotine. Exposure was at the ALI in a cloud chamber, which produced aerosols without heating. **(E.)** Modified experimental timeline used with EpiAirway™ tissue exposed at the ALI to aerosol with either a 24 hours (blue and cyan) or 48 hours (red and orange) recovery period, followed by viral pseudoparticle exposure at the ALI in a Cultex® exposure system. **(F.)** Flow cytometry data showing that protection against infection lasted at least 48 hrs after exposure to benzoic acid. Blue * and labels show comparison among the 24 hours groups. Red labels show comparisons among the 48 hours groups. **(G)** Flow cytometry data showing that BLU™ “Classic Tobacco” aerosol increased infection, but JUUL™ “Virginia Tobacco” aerosol did not. **(I)** Effect of power and chemical constituents in aerosols on infection of EpiAirway™ tissues. Exposures were done in a Cultex® system using a low power JUUL™ device that generated aerosol using 8 watts of power and a high power VOOPOO EC device that used 20 watts of power. All graphs are plotted as the mean ± standard deviation of three independent experiments. Box-Cox transformed data were analyzed using a one-way ANOVA followed by Tukey’s test to compare means. Black * = significantly different than the control. Color * = significantly different than comparison groups. * = p < 0.05, ** = p < 0.01, *** = p < 0.001, ****p < 0.0001. N.S. = not significant.

ALI exposure to aerosols containing PG/VG or PG/VG + nicotine significantly increased viral pseudoparticle infection of tissues **(Figure 2B)**. In contrast, JUUL™ aerosols caused a slight but significant decrease in infection. Tissues exposed to JUUL™ aerosols had significantly fewer infected cells than the PG/VG or PG/VG with 60 mg/mL nicotine groups. These data indicate that authentic JUUL™ aerosols negated the effect of PG/VG or PG/VG with 60 mg/mL nicotine on infection.

### Benzoic acid in EC aerosols decreased infection of EpiAirway™ tissues by viral pseudoparticles

Unlike PG/VG or PG/VG with nicotine, JUUL™ “Virginia Tobacco” fluid contains ∼44.8 mg/mL of benzoic acid and several flavor chemicals, which have been identified and quantified previously [27, 28]. The data in **Figure 2B** suggest that other chemicals in JUUL™ aerosols nullified the effect of PG/VG and nicotine on infection. To identify the chemical(s) that reduced infectivity, tissues were exposed to EC aerosols generated from: (1) PG/VG, (2) PG/VG with flavor chemicals (2,3,5 TMP, ethyl maltol, corylone and ethyl lactate = PG/VG+FLV), (3) PG/VG with 40 mg/mL of benzoic acid (PG/VG+BA), (4) PG/VG with flavor chemicals and benzoic acid (PG/VG+FLV+BA), (5) PG/VG with flavor chemicals, benzoic acid and 60 mg/mL nicotine (PG/VG+FLV+BA+NIC), and (6) JUUL™ “Virginia Tobacco” (**Figure 2C**). Flavor chemicals were used at the concentrations found in JUUL™ “Virginia Tobacco” [28]. Viral pseudoparticle infection was increased only in tissues exposed to PG/VG and PG/VG+FLV. All aerosols containing benzoic acid had infection equivalent to the clean air control (**Figure 2C**).

### Aerosols made with pure benzoic acid did not alter viral infection of the EpiAirway™ tissue

EC aerosols containing benzoic acid mitigated the effect of nicotine and PG/VG on viral pseudoparticle infection. We next tested pure aerosolized benzoic acid in the cloud chamber to determine if it alone alters infection by viral pseudoparticles. Nicotine aerosol increased viral pseudoparticle infection in EpiAirway™ (positive control), while benzoic acid alone had no effect **(Figure 2D)**.

### The protective effect of benzoic acid-containing aerosols lasted at least 48 hours after EC exposure in the Cultex® system

In our earlier experiments (**Figure 2B-2D**), tissues were allowed to recover for 24-hours before inoculation with viral pseudoparticles. To determine if benzoic acid in EC aerosol can provide protection against infection for a longer period of time, the recovery period was extended to 48 hours, as shown in the experimental timeline in **Figure 2E**.

PG/VG aerosol increased infection in the 24 and 48 hour recovery groups compared to the clean air controls. However, aerosols containing benzoic acid significantly reduced PG/VG-induced infection in both the 24- and 48-hour recovery groups **(Figure 2F)**. These results show that benzoic acid containing aerosols continue to protect EpiAirway™ tissues from infection at least 48 hours after the exposure had stopped.

### BLU™ disposable EC aerosol increased viral pseudoparticle infection, while JUUL™ aerosol with benzoic acid did not

BLU™ “Classic Tobacco” is a popular first-generation EC brand that has been marketed for many years and is still available. BLU™ disposable EC fluid contains 24 mg/mL nicotine and lacks benzoic acid, unlike JUUL™ EC fluid. Consistent with the previous infection results **(Figure 2B, 2C)**, EpiAirway tissues exposed to BLU™ aerosols showed an increase in viral infection **(Figure 2G)**. These data further support the idea that benzoic acid in EC aerosols reduces viral susceptibility and further show that EC aerosols that do not have benzoic acid increase infectivity.

### Comparison of a high and low power EC on viral pseudoparticle infection

JUUL™ ECs produce aerosols at relatively low power (8W-LP) [29], in contrast to third generation products that have variable and often higher powers [30]. VOOPOO, which is a popular high-power EC (20W-HP), produces a larger aerosol volume/puff that often increases aerosol complexity [31, 32]. PG/VG aerosols produced from both the low and high-power device increased viral pseudoparticle infection relative to the clean air control. The aerosols made with the high-power device produced a slightly higher percent of infection with a lower p-value than the low power device, but the groups were not significantly different. Infectivity in groups containing benzoic acid was equivalent to or slightly lower than the clean air control (**Figure 2H**). This protection was slightly less effective, in the benzoic acid aerosol generated in the high-power device.

### EC aerosols with benzoic acid reduced TMPRSS2 activity and prevented the increase in infection caused by PG/VG and nicotine

We next examined the effects of benzoic acid on ACE2 levels and TMPRSS2 activity. All EC aerosols tested increased ACE2 levels in the EpiAirway™ tissues compared to the clean air control (**Figure 3A, 3B**), although the increase in the BLU™ aerosol group was not significant (*p*=0.14). The inclusion of benzoic acid did not prevent this increase (**Figure 3B**), and benzoic acid by itself did not cause the increase when tested in the cloud chamber **(Figure 3C**).

**Figure 3.**
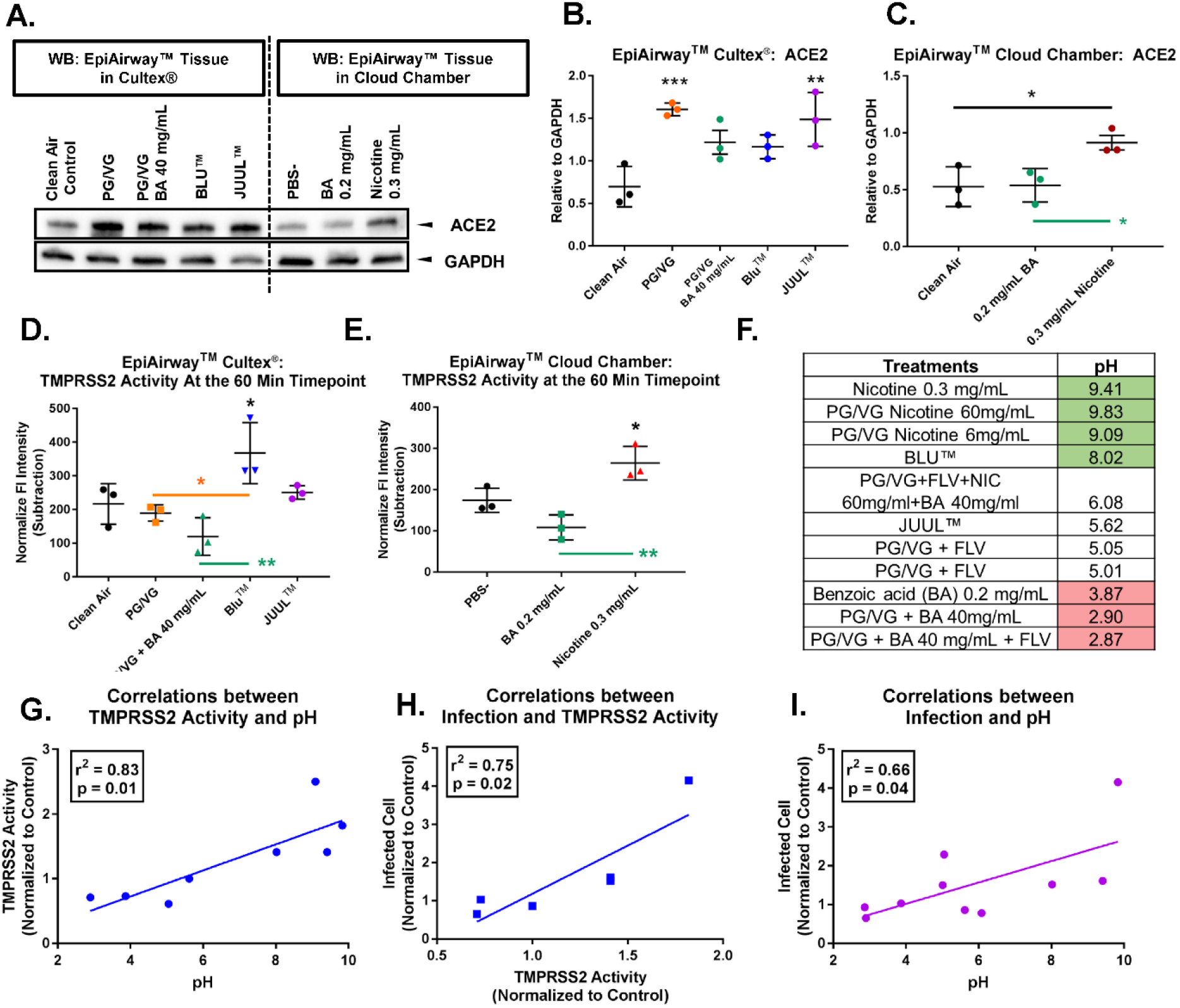
Benzoic acid in EC aerosol nullified PG/VG or nicotine enhanced infection in the EpiAirway™ tissue by decreasing TMPRSS2 activity. **(A - C)** The effect of aerosol exposures on ACE2 levels following exposure at the ALI in the Cultex® system **(A, B)** or Cloud Chamber **(A, C)**. Western blots were horizontally cropped from the original blot. (**D - E**) The effect of aerosol exposures on TMPRSS2 activity in the Cultex® system **(D)** or Cloud Chamber **(E)**. Graphs show activity at the 60-minute timepoint. **(F)** The table shows the pH of different treatment solutions and e-liquids before aerosolized exposure in the cloud chamber or Cultex® system. **(G - I)** Linear regression analysis comparing: **(G)** TMPRSS2 activity and pH, **(E)** infection and TMPRSS2 activity, and **(F)** infection and pH. Pairwise-Pearson correlation was used to test correlation and significance for each group. Data are plotted as the mean ± standard deviation of three independent experiments. In **B, C, D** and **E**,. Box-Cox transformed data were analyzed using a one-way ANOVA followed by Tukey’s posthoc test. Black * = significantly differently than the control. Color * = significantly different groups. * = p < 0.05, ** = p < 0.01, *** = p < 0.001, **** = p < 0.0001.

TMPRSS2 activity assays revealed that tissues exposed to aerosols containing PG/VG or PG/VG + benzoic acid had significantly lower activity than the control (**Figure 3D, E2A**). In contrast, tissues exposed to BLU™ aerosol, which lack benzoic acid, had a significant increase in TMPRSS2 activity. Exposure to pure benzoic acid in the cloud chamber decreased TMPRSS2 activity, but not significantly, while nicotine alone increased its activity **(Figure 3E, E2B)**. Overall, tissues exposed to aerosols containing benzoic acid showed a decrease in the activity of TMPRRS2.

### pH of test solutions and correlations between pH, TMPRSS2 Activity, ACE2 Levels, and infection

To further investigate the relationship between pH and viral infection, the pH values of lab-made and commercial e-liquids were measured (**Figure 3F)**. Solutions with nicotine, but not benzoic acid, had a high pH ranging from 8.02 in BLU™ to 9.83 in PG/VG + 60 mg/mL nicotine. Addition of benzoic acid neutralized the solutions and brought the pH down to 6.08 in lab-made liquids or 5.62 in authentic JUUL™ “Virginia Tobacco” e-liquid. Solutions with benzoic acid but without nicotine had lower pH values ranging from 2.87 – 3.87.

Linear regression analyses were performed on pH, ACE2 levels, TMPRSS2 activity, and infection to determine which factors affected infection (**Figure 3G-3I, E3A-E3B)**. There were weak positive correlations for ACE2 levels and pH (r^2^=0.34; **Figure E3A**), and for ACE2 levels and infection (r^2^=0.26; **Figure E3B**), but neither were significant (p > 0.05). TMPRSS2 activity showed a strong and significant positive correlation to pH (r^2^=0.83, p=0.01; **Figure 3G**). The correlation between TMPRSS2 activity and infection was also strong, but not significant (r^2^=0.66), however, when an outlier PG/VG point was removed, the correlation was stronger and significant (r^2^=0.75, p=0.02; **Figure 3H)**. There was a significant and strong positive correlation between pH and infection (r^2^=0.65, p=0.04; **Figure 3I**). The correlation table summarizes the results of linear regression analysis **(Figure E3C)**.

To determine the pH optimum for TMPRSS2, TMPRSS2 activity in clean air exposed tissue lysate was tested with buffers at different pH values (5,7,8,9). Optimal activity was observed between 7-8 (**Figure E3D**).

Taken together, these data demonstrated that ACE2 concentration does not always correlate with infectibility and that factors, such as pH, influence TMPRSS2 activity, which can alter infection.

## Discussion

Our results, which are summarized in Figure 4, provide evidence that PG/VG and nicotine-containing aerosols dose-dependently increased infection of EpiAirway™ 3D organotypic tissues by SARS-CoV-2 viral pseudoparticles and that benzoic acid in JUUL™ aerosols mitigates the effect of PG/VG or nicotine. The decrease in infection by benzoic acid was strongly correlated with reduced TMPRRS2 activity, which is required for viral fusion to host cells after binding to the ACE2 receptor [11, 12]. The protection from PG/VG and nicotine-enhanced-infection that benzoic acid provided remained at least 48 hours after exposure stopped. BLU™ EC aerosols, which contain nicotine, but not benzoic acid, increased pseudoparticle infection, showing that infection of EpiAirway™ varies with EC brand and e-liquid content. Infection of EpiAirway™ tissues occurred when both low- and high-power ECs were used to create aerosols, showing that even the low powered JUUL™ battery when used with a third-party pod was sufficient to induce PG/VG or nicotine enhanced infection. TMPRSS2 activity was highly correlated with the pH of the aerosolized e-liquid, and infection was highly correlated with pH, with low pH values producing low levels of infection. These data are consistent with the conclusion that the relationship between EC use and SARS-CoV-2 infection depends on the e-liquid ingredients, which can either increase or protect against nicotine induced infection by SARS-CoV-2 pseudoparticles.

**Figure 4:**
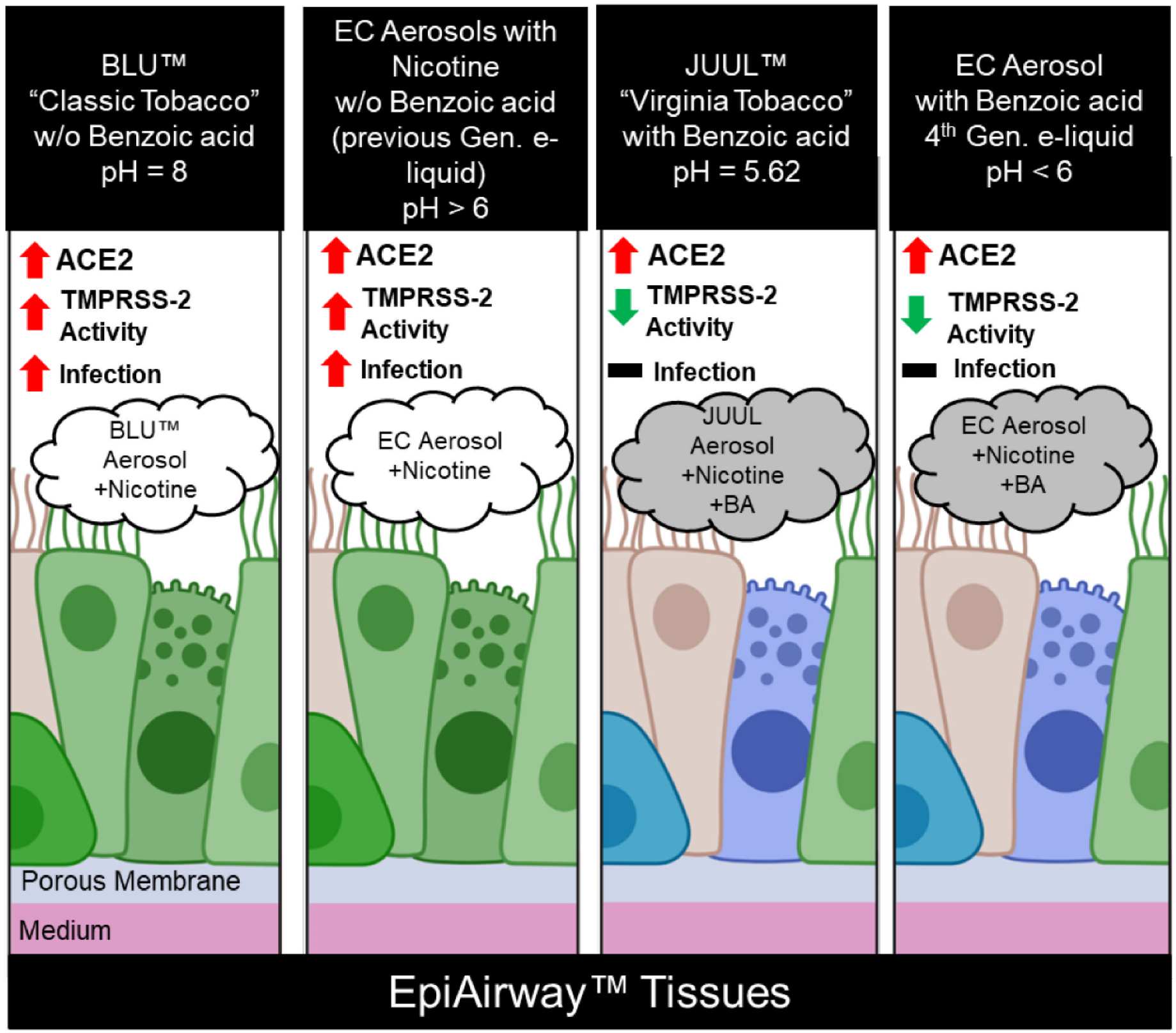
The effects of nicotine, benzoic acid, and pH on ACE2 levels, TMPRSS2 activity, and SARS-CoV-2 pseudoparticle infection. Summary of the responses of EpiAirway™ tissues to aerosols produced from authentic BLU™ “Classic Tobacco”, authentic JUUL™ “Virginia Tobacco”, mixtures (PG/VG plus nicotine, PG/VG plus nicotine and benzoic acid), and individual chemicals (PG/VG, nicotine, benzoic acid). All EC aerosols increased ACE2 levels in the EpiAirway™ tissue. However, the infectivity of the EpiAirway™ tissues by the SARS-CoV-2 pseudoparticles was not directly correlated with ACE2 levels. The pHs of the e-liquid modulated TMPRSS2 activity which was a critical factor that affected the infectivity by the SARS-CoV-2 viral pseudoparticles. E-liquids with nicotine, but not benzoic acid, such as BLU™ “Classic Tobacco”, had pHs >6, which elevated TMPRSS2 activity and enhanced infection. E-liquids with nicotine and benzoic acid, such as JUUL™ “Virginia Tobacco”, had lower pHs ranging from 5.62 – 6, which reduced TMPRSS2 activity and mitigated nicotine-enhanced infection.

We designed our Cultex® exposure protocol to simulate an acute EC exposure over 3 days. EC exposures vary among users and can range from 1 to 1,265 puffs/day with reported averages of 140 and 200 puffs/day [26, 33]. We chose 50 puffs/day as this is within the range that an EC user receives, although it is below several reported averages. The Cultex® system generates authentic EC aerosols, which include any chemicals present in the e-liquid plus reaction products and metals that are added during heating of the e-liquid [34-37]. The cloud chamber enabled further study of the effects observed in the Cultex® system through isolation of the chemicals. The combination of these two ALI exposure systems provides an innovative technology for understanding how aerosols affect the human respiratory system, and when used in conjunction with 3D EpiAirway™ models represent a powerful approach for assessing the effect of viruses on the respiratory system, a problem that cannot readily be addressed experimentally in human subjects.

In our previous study, we used monolayers of BEAS-2B cells in submerged cultures and ALI exposures [21]. Treatments that increased ACE2 (nicotine, nicotine plus PG/VG, and authentic JUUL™ aerosol) also increased viral pseudoparticle infection; however, the increase in the JUUL™ group was significantly less than in the nicotine only group, even though both groups were exposed to a similar concentration of nicotine [21]. It is likely that benzoic acid in JUUL™ aerosol suppressed nicotine-enhanced infection in our prior study, but the effect was not as great as in the current study. This difference is likely due to variations in the exposures. In the BEAS-2B study, cells in the Cultex® were exposed only once to 10 puffs of JUUL™ aerosol. In the current study, exposures were 50 puffs/day over a 3-day span. The latter exposure protocol was sufficient to completely prevent nicotine enhanced infection of EpiAirway™, while 10 puffs on one day only partially reduced the effect of nicotine on infection of BEAS-2B cells. Comparison of these studies shows that user topography, which affects nicotine exposure, could affect the outcome of experiments and the health effects that EC users experience.

A critical result in this study was the finding that authentic JUUL™ aerosols and mixtures containing benzoic acid kept SARS-CoV-2 pseudoparticle infection at or below clean air control levels. Although benzoic acid has been used for many years as a food preservative [38] and recently became a common additive in 4^th^ generation EC products [39], its effect on the respiratory epithelium is not well characterized. Our data showed that EpiAirway™ treated with EC aerosols containing benzoic acid caused infection results similar to the control group, while aerosols from BLU™, which does not have benzoic acid, elevated infection. Benzoic acid by itself did not affect infection (cloud chamber), but when combined with EC chemicals that normally increase infection, such as nicotine or PV/VG, benzoic acid held infection to control levels. These data clearly show that infection by SARS-CoV-2 is strongly influenced by the ingredients in e-liquids and inclusion of benzoic acid can reverse the increases in infection induced by nicotine and PG/VG.

The suppression of PG/VG or nicotine-enhanced infection by benzoic acid was correlated with EC-liquid pH and with TMPRSS2 activity. These data support the conclusion that benzoic acid lowers the pH at the ALI to a level that reduces TMPRSS2 activity, which in turn reduces infection. This conclusion is supported by our data showing that TMPRSS2 had optimal activity between pH 7-8, well above that in benzoic acid-containing solutions. Our optimum is in reasonable agreement with Shrimp et al. [24] who reported the pH optimum of isolated TMPRSS2 protein to be between 8-9. The differences in range for pH optima may be due to the use of cell lysates vs. isolated enzyme. In either case, TMPRSS2 has optimal activity at 7 or above, and our infection data show that aerosols produced from e-liquids with higher pH (i.e., those containing nicotine without benzoic acid or made from BLU™ EC) in most cases had higher levels of infection. Other types of viruses may also be susceptible to the low pH levels in benzoic acid containing e-liquids. For example, influenza A type virus undergoes an irreversible loss of activity at pH levels between 4.6 and 6.0 when tested in vitro [40].

The low pH aerosols containing benzoic acid may affect binding of the virus to its ACE2 receptor. The spike protein has three receptor binding domains (RBDs) that are cryptic at low pH and become exposed and available for binding as pH is elevated [11, 41]. Additionally, pH 7.5-9 was found the be an optimum pH for binding of ACE-2 and spike [42]. Low pH aerosols likely reduce the exposure of the cryptic RBDs and therefore also reduce the ability of the pseudoparticles to bind to ACE2. The pH of exhaled aerosol may also inactivate both influenza A and SARS-CoV-2 viruses in indoor environments [43], suggesting that the low pH of JUUL™ aerosols may be directly damaging to SARS-CoV-2 virus. This in combination with reduced TMPRSS2 activity are mechanisms that likely reduced viral infection in JUUL™ and other aerosols containing benzoic acid.

The suppression of nicotine- or PG/VG-induced infection by benzoic acid was sustained 48 hours after ALI exposure, suggesting that benzoic acid may have additional roles in reducing infection. While pH may be lowered during inhalation of benzoic acid-containing aerosols during vaping, it likely returns to normal after vaping has stopped, and this probably does not take 48 hours to occur. Our study focused on the infection step of COVID-19. It is possible that benzoic acid also affects later stages in pathogenesis. *In silico* models have shown that benzoic acid and its derivatives can interact with proteases involved in SARS-CoV-2 replication and reduced their functions [44].

In contrast to nicotine-containing aerosols, aerosols produced from PG/VG alone had a low pH (5), yet increased infection. This may be due to PG’s ability to stabilize protein interactions and to increase transepithelial permeability. PG can stabilize small molecules and proteins and extend their reactivity for a long period [45]. Prior to SARS-CoV-2 entry, spike protein is cleaved by furin at the boundary of the S1 and S2 subunits. This modification increases instability of spike and can cause the S1 subunit to be shed from spike, leading to a premature change to the post-fusion conformation of spike, which would make it non-functional [11]. Exposure to PG/VG aerosol may help stabilize the tenuous association between the S1-S2 subunits, increasing the chance of infection. In support of this idea, the SARS-CoV-2 D614G strain acquired a mutation that stabilized the S1-S2 association and increased its infectability [46]. PG/VG e-liquid treatments also rapidly decrease transepithelial electrical resistance, weaken cellular tight junctions, and increase transepithelial permeability [47]. Such weakening of cellular-tight junctions could allow the virus to penetrate deeper into the tissue, causing more infection.

Our finding that nicotine increased ACE2 agrees with prior studies [15-24]. To the best of our knowledge, our data are the first to show that PG/VG aerosol can increase ACE2, an important finding since these solvents are the dominant chemicals in e-liquids [2, 48]. We also showed that both nicotine and PG/VG increase ACE2 when present in authentic JUUL™ aerosols made using the Cultex® system. The parallel exposures in a cloud chamber and Cultex® thus enabled authentic aerosol to be evaluated and the chemicals responsible for ACE2 elevation to be identified. Elevation of ACE2 by nicotine is generally considered an indicator that viral infection would increase [15-18, 49]. While our data in part support this conclusion (e.g., PG/VG and nicotine elevated ACE2 and pseudoparticle infection), increases in ACE2 were not always correlated with increased infection. For authentic JUUL™ aerosols and mixtures containing benzoic acid, ACE2 was usually significantly elevated, but infection was at or below clean air control levels. Thus, caution should be used when interpreting ACE2 levels, as they do not always correlate with infection.

Increases in PG/VG-induced infection of EpiAirway™ occurred with both the VOOPOO (20 watt) and JUUL™ (8 watt) ECs. While the VOOPOO’s infection was higher than that of the JUUL™ with the third-party pod, there was not a significant difference between the groups, showing that even low powered ECs can produce the PG/VG-induced increase in infection as effectively as the higher powered VOOPOO. In both cases, benzoic acid prevented the PG/VG-induced increases in infection.

Most work on the relationship between tobacco product use and COVID-19 has been done with cigarette smokers and tobacco cigarettes. Several recent meta-analyses found that cigarette smokers have a higher susceptibility to SARS-CoV-2 infection than non-smokers, and following infection, patients with smoking histories were more likely to develop severe COVID-19 symptoms leading to hospitalization and death [50-52]. Nevertheless, the relationship between smoking and COVID-19 remains unresolved and some investigators have proposed that nicotine, from tobacco smoke, protects against SARS-CoV-2 infection [53,54].

Similarly, there is some discrepancy in the EC literature regarding the influence of EC use to SARS-CoV-2 susceptibility [15-23]. Our study clearly shows that the presence of benzoic acid in e-liquids mitigates the enhanced infection caused by nicotine. Because prior EC studies with humans have not segregated EC users based on whether they used acid containing ECs, the data are likely noisy, which may account for some of the contradictory reports regarding the correlation between EC use and SARS-CoV-2 infection. In two clinical studies [22, 23], all EC user data were collected into a single group without regard to the EC products used. Based on our data, it would be interesting, if possible, to reanalyze these clinical data and separate the EC users into subgroups based on their EC products (acid containing or not). Perhaps, another conclusion could be extracted and could explain some conflicting information surrounding EC use and SARS-CoV-2 susceptibility.

In summary, our data address the question: “Does EC use increase or decrease the likelihood of SARS-CoV-2 infection?”. The answer is EC use can do either depending on the ingredients in the e-liquid that is aerosolized. EC aerosols, with or without nicotine and with or without benzoic acid, increased ACE2 in the EpiAirway™ tissues, and this could facilitate infection by SARS-CoV-2. However, TMPRSS2 activity, which is seldom monitored, was decreased at the acidic pH of EC aerosols containing benzoic acid and this held infection to clean air control levels. Many 4^th^ generation EC products use benzoic or other acids that lower aerosol pH, and these are also likely to mitigate the enhanced infection. Given the diversity of e-liquids that have been sold in the past and currently, it is not surprising that prior studies have drawn different conclusions regarding the relationship between vaping and COVID-19 [15-23]. Overall, these data demonstrate that individual ingredients (nicotine, PG/VG, and benzoic acid) can influence SARS-CoV-2 infection of human bronchial epithelial tissues and that benzoic acid, if present, will override the increased infection produced by nicotine and PG/VG. Future studies involving organotypic cultures, animals, or humans should separately evaluate groups with and without e-liquids containing benzoic or other acids. If low and high pH e-liquids are not studied separately, the data will likely be unclear and significant effects of vaping will be missed. We do not advocate acid containing EC products as a prophylactic for avoiding COVID-19; however, if already vaping, then using an acid containing product may help reduce viral infection. It is important to be aware that the long-term effects of inhaling benzoic or other acids in the context of EC aerosol is not known and may itself have adverse health consequences.

### Limitations of the Study

We examined one 4^th^ generation product (JUUL™ “Virginia Tobacco”). It will be important to determine if aerosols produced from other 4^th^ generation ECs that have benzoic and other acids can offset the effects of nicotine and PG/VG. The EpiAirway™ model provides an *in-vitro* analysis technology that closely resembles *in-vivo* exposure in humans. Because standard MatTek EpiAirway™ uses primary epithelium from a single donor, it would be informative to test 3D cultures from other human donors. The effect of acid pH on infection lasted at least 48 hours. In the future, other acids and longer intervals after exposure could be tested. Finally, during heating of e-liquids, metals can be added to the aerosols and reaction products can form from the solvents and flavor chemicals [33-36]. It is not yet known how these chemicals affect SARS-CoV-2 infection. While the flavor chemicals we tested did not later infection percentage, it is possible that other flavor chemicals can have an effect.

## Author Contributions

Project administration and funding acquisition, P.T.; Conceptualization, R.P. and P.T.; Investigation R.P.; Sample preparation, data collection, and data processing, R.P., M.W., A.S., and T.M.; Data Interpreted, R.P. and P.T.; Writing the original draft, R.P. and P.T.; Reviewing and Editing, R.P., P.T., and T.M.

## Acknowledgements

We thank Dr. James Pankow, Wentai Lou, Kevin McWhirter from Portland State University for quantifying the concentrations of benzoic acid deposited in the ALI exposure systems. We would like to acknowledge the contribution to Jack Ona and Claudia Osuna for assisting in viral pseudoparticle productions. We would like to thank the Stem Cell Core at UCR for providing some of the equipment that was used in this investigation. Diagram in Figure 5 was created with BioRender.com.

## Supplementary Materials and Methods

### HEK 293T ^ACE2^ Cell Culture and Maintenance

HEK 293T cells (ATCC, Manassas, VA, USA), and HEK 293T cells, which over expressed ACE2 (293T^ACE2^, BEI resource, Manassas, VA, USA; NR-52511) were cultured in DMEM high glucose medium supplemented with 10% fetal bovine serum (FBS; ATCC, Manassas, VA, USA). At 80-90% confluency, cells were washed and detached as described above. Cells were seeded at 3,000 cells/cm^2^ and incubated in a 37°C, 5% CO_2_, 95% relative humidity incubator.

### EpiAirway™ Tissue Maintenance

EpiAirway™ primary cell tissues cultured at the ALI have beating ciliated cells, mucus producing goblet cells, and keratin-5-positive basal cells. Upon arrival, EpiAirway^™^ inserts from MatTek Corporation (Ashland, MA, USA) were transferred from the agarose shipping plate to a 12-well culture plate pre-filled with AIR-100-ASY medium in basal-lateral compartments. These tissues were acclimated and stored in an incubator with 37°C, 5% CO_2_, and 95% relative humidity for 24 hours prior to any experiments. Basal lateral medium was replaced every day. Prior to aerosol exposure, the apical side of the EpiAirway^™^ was washed with Dulbecco’s phosphate buffered saline with calcium and magnesium (DPBS+; Lonza, Walkersville, MD, USA).

### Reagents and Procedure for Compounding Lab-Made EC Refill fluids

Refill fluids contained propylene glycol (PG; Thermofisher, Tustin CA, USA) and vegetable glycerin (VG; Thermofisher, Tustin CA, USA) made at a 30 PG/70 VG ratio. Liquid (-)- nicotine (Sigma-Aldrich, St. Louis, MO, USA) was added to PG/VG to obtain either 6 or 60 mg/mL of nicotine. In some refill fluids, benzoic acid was dissolved in heated VG and allowed to cool at room temperature; PG was then added to make a 30 PG/70VG ratio containing 40 mg/mL of benzoic acid (Sigma-Aldrich, St. Louis, MO, USA), which is similar to the reported concentration [1]. To replicate JUUL™ “Virginia Tobacco” refill fluid, the VG with benzoic acid solution was mixed with PG containing 2,3,5 TMP, ethyl maltol, corylone, and ethyl lactate (Sigma-Aldrich, St. Louis, MO, USA) to give the following final concentrations: 0.027 mg/mL 2,3,5 TMP, 0.033 mg/mL ethyl maltol, 0.036 mg/mL corylone, 0.125 mg/mL ethyl lactate, 40 mg/mL benzoic acid, and 60 mg/mL nicotine in a 30 PG/70 VG solvent, as reported in previous publications [1,2].

### Source of EC Refill fluids, EC Devices, and Pods

Blu™ ECs, JUUL™ ECs, Drag S VOOPOO ECs, VOOPOO “PnP” pods and coils, JUUL™ “Virginia Tobacco” pods, and blank JUUL™ compatible pods (OVNS JC01 pods with 1.5Ω ohm ceramic wick) were purchased online from third party vendors, at a local gas station, and at grocery stores. JUUL™ “Virginia Tobacco” refill fluids were extracted from pods by removing the cover cap, rubber stopper, and directly collecting the refill fluids.

### Aerosol exposures at the ALI in the VITROCELL® cloud chamber

Monolayers of EpiAirway™ tissues cultured on 12 well Transwell® inserts were placed into a VITROCELL® cloud chamber (VITROCELL®, Walkirch, Germany) for an ALI exposure to various chemical aerosols that were generated without heating. Prior to exposures, pre-warmed culture medium was added into wells in the exposure chamber and allowed to equilibrate to 37 ºC. Stock nicotine was diluted with PBS- to make exposure solutions with final concentrations either 0.3 mg/mL nicotine. Benzoic acid flakes were dissolved in warm water and diluted with PBS- to make an exposure solution with a final concentration of 0.2 mg/mL benzoic acid. For each exposure, 200 μL of exposure solution were added into a VITROCELL® nebulizer to generate a uniform aerosol with a flow rate of 200 μL/min without using heat. Control cells were exposed to PBS-aerosols.

EpiAirway™ tissues were exposed to 5 puffs of PBS-, nicotine (0.3 mg/mL), or benzoic acid (0.2 mg/mL) aerosol. The aerosols were allowed to settle onto cells until the chamber becomes visually clear, which takes 3 mins. Between puffs, the cloud chamber was allowed to vent for 1 minute. After the exposure, the cells were returned to the incubator to recover for 24 hrs.

### EC aerosol exposure at the ALI in the Cultex® RFS compact exposure system

A Cultex® RFS compact exposure module (Cultex Laboratories GmbH, Hannover, Germany) was used to expose cells or tissue cultures to humified sterile air (clean air control) or EC aerosols generated from EC devices. Prior to each exposure, cells or tissues samples were placed into the exposure chambers, which contained culture medium and were regulated at a temperature at 37 ºC.

The Cultex® exposure system consisted of a sampling module and an aerosol guiding module. The sampling module used a custom designed EC smoking robot (RTI International, NC, USA) that draws filtered air in the biosafety cabinet or EC aerosol from the EC device into a 200 mL syringe. After collecting 55 mL of either filtered air or aerosols, the Cultex® system then dispense the sampled air or aerosols into the aerosol guiding module, which mixes either the filtered air or EC aerosols with humified zero air (1 L/min). This mixture diluted the aerosol and generated a uniform flow before aerosols were directly distributed onto each tissue sample. Exposure mixtures were allowed to settle onto cells for 5 second and then vented out the exposure chamber at a flow rate of 5 mL/min, generated by a mass flow controller (Boekhorst, Bethlehem, PA, USA), and finally dispensed into a waste container.

Each exposure consisted of 55 mL of filtered air or EC aerosol. Puffs lasted 4 seconds and were spaced 30 seconds apart. EpiAirway™ 3D tissues were exposed to 50 puffs of air or EC aerosols/day for 3 days. Between each exposure day, tissues were returned to the incubator and after the last exposure, tissues were allowed to recover in the incubator for 24 hrs prior to analyses.

### Immunofluorescence of EpiAirway™ Tissues

After EC aerosol exposures and recovery, EpiAirway™ tissues were fixed in 4% paraformaldehyde (PFA) at 4°C for 20 hrs, then the apical and basal surfaces were washed with DPBS several times. Fixed tissues were stored in cold DPBS solution and shipped to MatTek for histology. The samples were returned to us as unstained histology sample slides, which were rehydrated through a graded series of ethanol washes, until reaching pure nano water. Antigen retrieval was performed using a 10 mM citrate buffer (pH 6.0) in a 95 ºC water bath for 20 mins. Permeabilization, blocking, and primary and secondary antibody incubation steps were the same as for immunocytochemistry. Samples slides were mounted using Vectashield with DAPI (Vectashield, San Francisco, CA, USA) and glass cover slips. Fluorescent cells were imaged with a Nikon Eclipse Ti inverted microscope (Nikon Instrument, Melville, NY, USA) using a 60X objective, and images were captured using a high-resolution Andor Zyla VSC-04941 camera (Andor, Belfast, UK). Antibodies used were anti-ACE2 (1:200; R&D system, Minneapolis, MN, USA), anti-TMPRSS2 (1:200; Santa Cruz, Dallas, TX, USA), anti-acetylate α-tubulin conjugated to Alexa fluor-594 (1:200; Santa Cruz, Dallas, TX, USA), anti-MUC5AC conjugated to Alexa fluor-594 (1:100; Santa Cruz, Dallas, TX, USA), and anti-keratin 5 (1:200, Biolegend, San Diego, CA, USA). Secondary antibodies were Alexa fluor-488 or Alexa fluor-594 (Thermofisher, Tustin CA, USA).

### Lysate Preparation for Western Blot and Proteolytic Assay

After exposure at the ALI, RIPA buffer with or without PMSF protease inhibitor (ChemCruz Biochemical, Dallas, TX, USA) was used to lyse cells and tissues. The lysates prepared for Western blot use RIPA buffer with protease inhibitor, while lysates prepared for proteolytic assay used RIPA buffer without a protease inhibitor. The cell lysates were vortexed every 10 mins for 30 mins, pipetted through 23-gauge needles several times, then centrifuged at 3,000 x g for 5 mins at 4°C. The lysate protein was quantified using the Pierce BCA assay kit (Thermo Scientific, Waltham, MA, USA). Each Western blot used 20 μg of protein, and each proteolytic assay used 5 μg of protein.

### Western Blot

Following the lysate preparation step, denaturing buffer (β-mercaptoethanol and Laemeli buffer, 1:10) was added to each Western blot lysate at a 1:4 ratio. The buffer/lysate mixtures were heated at 95°C for 2 mins, then loaded onto an SDS gel (BioRad, Carlbad, CA, USA) for electrophoretic separation of proteins (120V for 1-2 hrs), and then transferred onto a PVDF membrane (BioRad; 200mA overnight at 4°C). After transfer, the membrane was cut horizontally, either under or above the expected location of the protein of interest based on its molecular weight (kDa). Membranes were blocked with 5% milk in TBS-T (TBS with 1%Tween-100) buffer for 2 hrs. and incubated overnight at 4°C with antibodies against anti-ACE2 (1:400; R&D systems, Minneapolis, MN, USA), anti-TMPRSS2 (1:1000; Santa Cruz, Dallas, TX, USA), anti-GADPH (1:2000; Cell Signaling Technology, Danvers, MA, USA). Membranes were washed for 30 mins in TBS-T, then incubated in an HRP conjugated secondary antibody (1:1000; Santa Cruz, Dallas, TX, USA or Cell Signaling Technology, Danvers, MA, USA) for 2 hrs. at room temperature. Membranes were developed using BioRad Clarity^™^ Western ECL Substrate reagent (BioRad, Carlbad, CA, USA) in a BioRad ChemiDoc^™^ Imaging Systems (BioRad, Carlbad, CA, USA), which determined the optimal exposure time for each protein (exposure times were ACE2 27ms; TMPRSS2 24ms; GADPH 4ms).

### Spike Viral Pseudoparticle Production

Pseudoparticle production was performed as outlined in Crawford et al., 2020 [3]. In brief, HEK293T cells were plated with antibiotic-free medium at a density of 7 × 10^6/^/T75 flask and transfected with Lipofectamine3000 (Thermo Fisher Sci, Waltham, MA, USA) using a total of 15 μg of BEI lentiviral plasmids (BEI resources, Manassas, VA, USA).

1. NR-52520: pHAGE2-CMV-ZsGreen-W7.5 μg
2. NR-52517: HDM-Hgpm2, 1.65 μg
3. NR-52519: pRC-CMV Rev1b, 1.65 μg
4. NR-52518: HDM-tat1b, 1.65 μg
5. NR-52314: pHDM-SARS-CoV-2 Spike, 2.55 μg.

60 μL of Lipofectamine and 20 μL of P3000 reagent was used following the manufacturer’s protocol. After overnight incubation, fresh medium supplemented with 1% bovine serum albumin (BSA; Sigma Aldrich, St. Louis, MO, USA) was added to the cells. Fluorescence microscopy was used to visually inspect transfection efficiency and the expression of the ZsGreen. The cell culture medium was collected 48 hours after transfection, centrifuged, and the supernatant was filtered with a 0.45 μm syringe filter. The filtered supernatant was mixed with 5X Polyethlyene glycol (Abcam, Cambridge, UK) and precipitated overnight at 4°C. The lentivirus was collected by centrifugation and resuspension of the pellet in Viral Re-suspension Solution (Abcam, Cambridge, UK). The virus aliquots were stored at -80°C. Prior to infection experiments, the transduction efficiency of each viral batch was determined by infecting 293T^ACE2^ cells and quantifying the number of infected cells with flow cytometry.

### Flow cytometry

Prior to flow cytometry, fluorescence microscopy was used to validate the expression of ZsGreen signal. All samples were pipetted several times to generate single cell suspensions and passed through a 35 μm filter of a Falcon^™^ Round-Bottom 5 mL polystyrene test tube (Fisher Scientific, Tustin, CA, USA). Forward-Scatter-Height (FSC-H) and Forward-Scatter-Area (FSC-A) were used to generate a gate to select single cell for each sample. Side-Scatter-Area (SSC-A) and FSC-A were used to generate a gate to exclude small debris. Non-fluorescent mock infected cells were used to generate a gate to quantify fluorescence signal of infected cells. Final results were represented in percent of fluorescent infected cells in each sample.

### Measuring pH of lab made, BLU™ and JUUL™ “Virginia Tobacco” e-liquid

The pH meter (Thermofisher, Tustin, CA, USA) was calibrated with solutions at pH 4, 7, and 10 before measuring the pH of each diluted e-liquid sample. Lab made, Blu™ and JUUL™ e-liquid were diluted at a 1:10 ratio in ultrapure water, then the pH of the diluted sample was measured. For each session of measurement, the pH electrode is fully submerged in the sample. Between each sample, the pH electrode was rinsed with ultra-pure water and patted dry before measuring new samples.

### Correlation plots

ACE2 concentration, TMPRSS2 activity, and infection data were normalized to their respective control groups. The normalized data were then used to calculate correlations between each group in Minitab statistic software (Minitab, State College, PA, USA). Pairwise Pearson correlation was used to test correlation and significance between each group, and GraphPad Prism 7 (GraphPad, San Diego, CA, USA) was used to plot the linear regression graphs.

### Statistical Analysis

In all experiments, three independent EpiAirway tissues were used. Statistical analyses of all data were done with Minitab Statistics Software (Minitab, State College, PA. USA). If data satisfied the assumptions of ANOVA (homogeneity of variances and normal distribution), they were analyzed using a one-way ANOVA. If the assumptions were not met, data were subjected to a Box-Cox transformation, and retested to verify they satisfied the assumptions of ANOVA. When the ANOVA detected significance (p = 0.05), Tukey’s postdoc test was used to compare group means. In the TMPRSS2 activity assays, two-way ANOVAs were used to study the effects of treatment and time. Means were considered significantly different for p > 0.05. Data were plotted using GraphPad Prism 7 software (GraphPad, San Diego, CA, USA)

---

**Figure E1.**
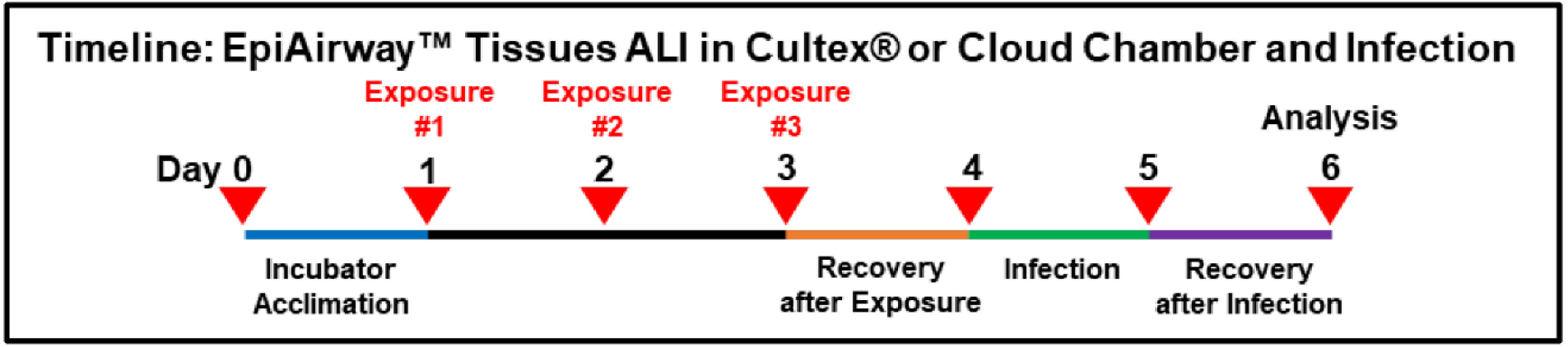
Timeline showing experimental design used with EpiAirway™ tissue exposed at the ALI to aerosols in a Cultex™ exposure system or cloud chamber, then infected with viral pseudoparticles for 24 hours.

**Figure E2.**
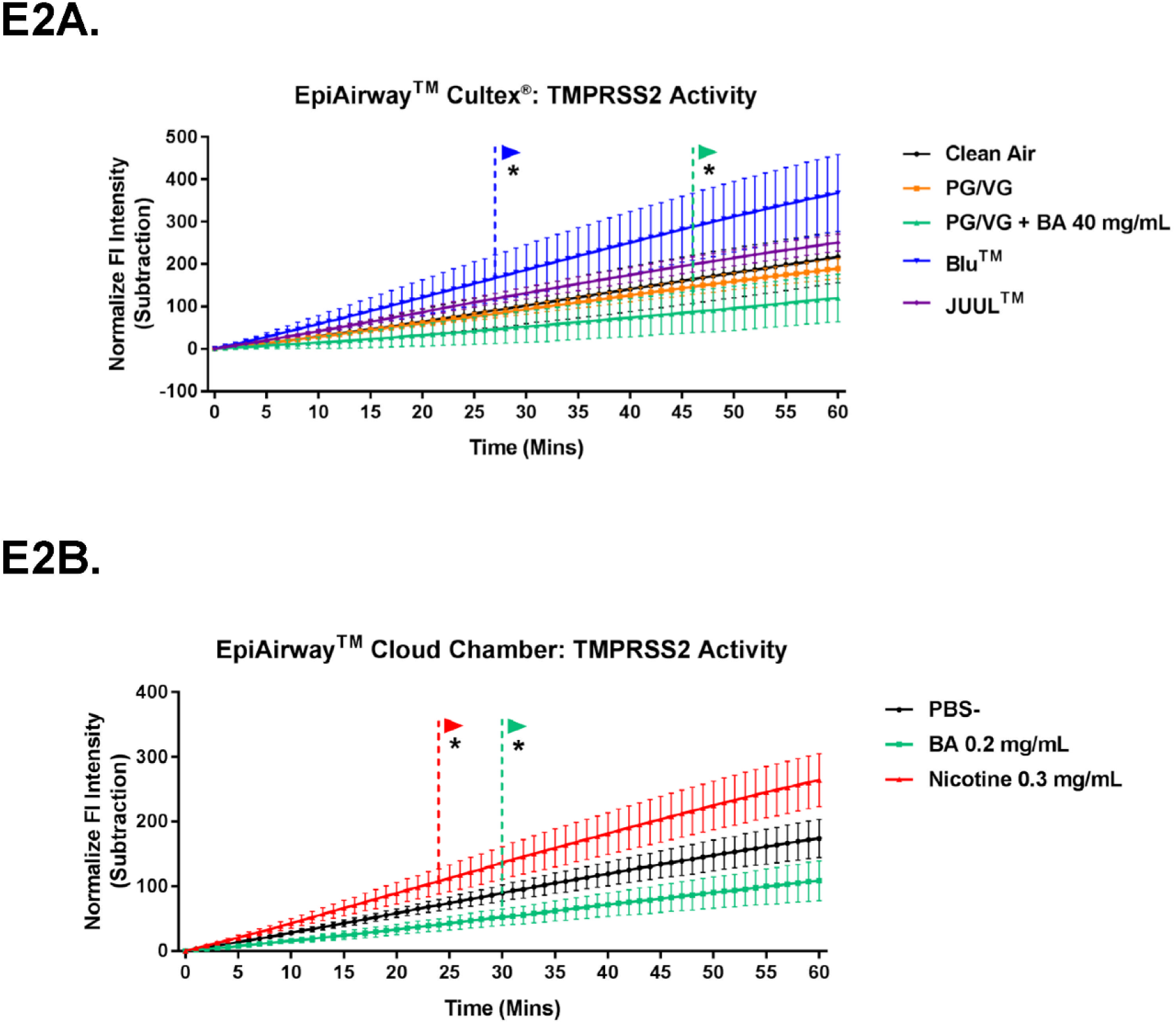
Benzoic Acid in EC Aerosol Nullified PG/VG or Nicotine Enhanced Infection in the EpiAirway™ Tissue by Decreasing TMPRSS2. TMPRSS2 activity assays showing effects of EC aerosols on the EpiAirway™ tissues in the Cultex® system (**E2A**) or in the cloud chamber (**E2B**). Both graphs show activity over 60 minutes. * indicate when values became significantly different than the control (*p* ranged from 0.05 to 0.0001). Data were analyzed using a two-way ANOVA followed by Dunnett’s posthoc test to compare means to the exposure control

**Figure E3.**
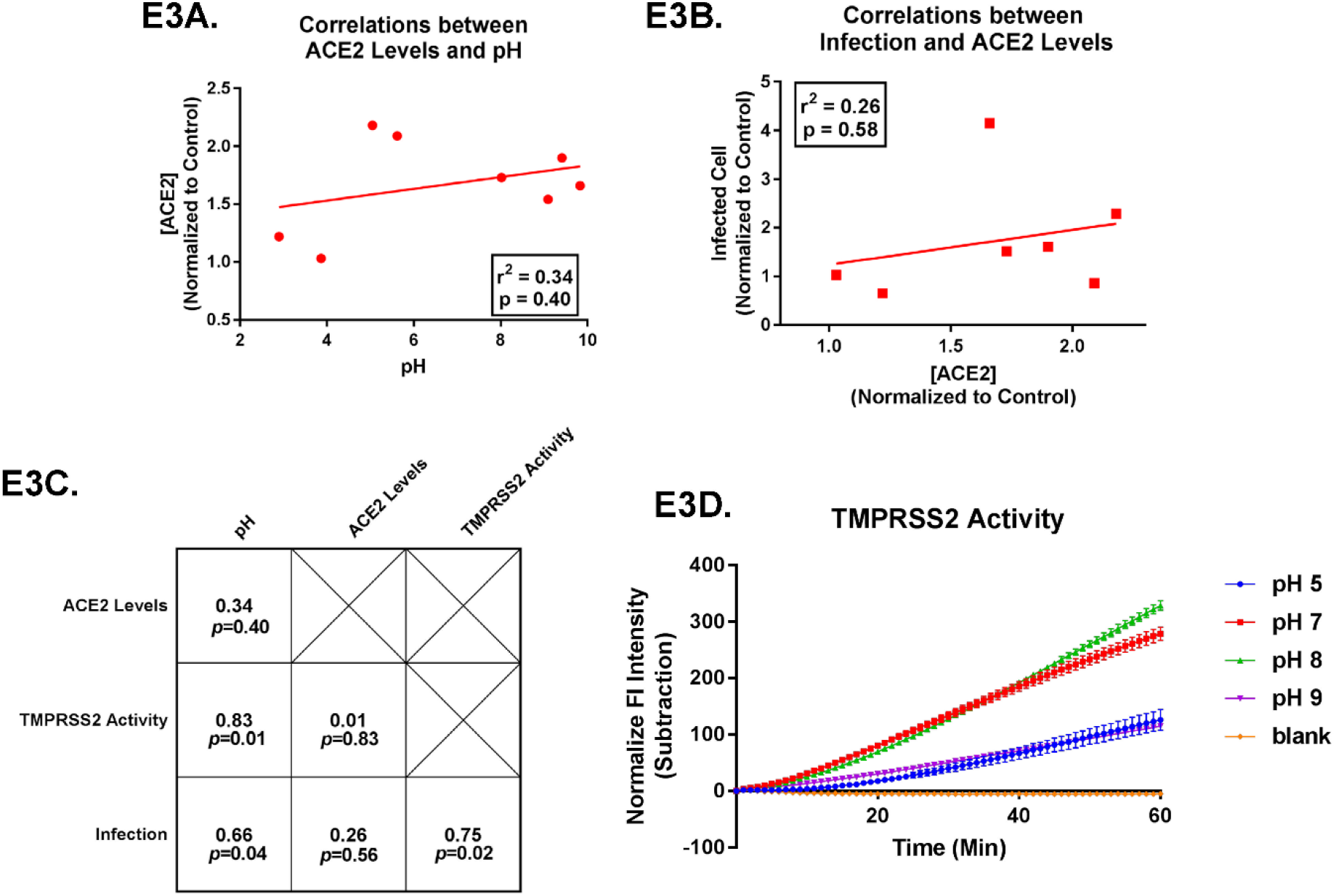
The correlation between e-liquid pH, ACE2, TMPRSS2 activity, and Infection. **(E3A-E3B**) Linear regression analysis comparing: **(E3A)** ACE2 concentration and pH and **(E3B)** infection and ACE2 concentration **(E3C)** The correlation table summarizes each relationship of e-liquid, ACE2, TMPRSS2, and infection. R^2^ values show for each interaction. The color gradient indicates the strength of each interaction from gray (r^2^=0 to red (r^2^=1). **(E3D)** Graphs show TMPRSS2 activity of cell lysates in buffers with different pHs over 60 minutes.

## References

1. Trtchounian A, Talbot P. Electronic nicotine delivery systems: Is there a need for regulation? Tobacco Control. 2011;20(1):47–52.

2. Omaiye EE, Luo W, McWhirter KJ, Pankow JF, Talbot P. Disposable puff bar electronic cigarettes: chemical composition and toxicity of e-liquids and a synthetic coolant. Chemical Research in Toxicology. 2022;25(8), 1344–58.

3. Pisinger C, Døssing M. A systematic review of health effects of electronic cigarettes. Preventive Medicine. 2014;69:248–60.

4. Sakamaki-Ching S, Williams M, Hua M, Li J, Bates SM, Robinson AN, Lyons TW, Goniewicz ML, Talbot P. Correlation between biomarkers of exposure, effect and potential harm in the urine of electronic cigarette users. BMJ Open Respiratory Research. 2020;7(1):e000452.

5. Hua M, Sadah S, Hristidis V, Talbot P. Health effects associated with electronic cigarette use: automated mining of online forums. Journal of Medical Internet Research. 2020;22(1):e15684.

6. Pozuelos GL, Kagda M, Rubin MA, Goniewicz ML, Girke T, Talbot P. Transcriptomic evidence that switching from tobacco to electronic cigarettes does not reverse damage to the respiratory epithelium. Toxics. 2022;10(7): 370.

7. Gotts JE, Jordt S-E., McConnell R, Tarran R. What are the respiratory effects of e-cigarettes? BMJ. 2010;l5275.

8. Sussan TE, Gajghate S, Thimmulappa RK, Ma J, Kim JH, Sudini K, et al. Exposure to electronic cigarettes impairs pulmonary anti-bacterial and anti-viral defenses in a mouse model. PLoS One. 2015;10(2):e0116861.

9. Reidel B, Radicioni G, Clapp PW, Ford AA, Abdelwahab S, Rebuli ME, et al. E-cigarette use causes a unique innate immune response in the lung, involving increased neutrophilic activation and altered mucin secretion. Am J Respir Crit Care Med. 2018; 197(4):492–501.

10. Goldenson NI, Leventhal AM, Stone MD, McConnell RS, Barrington-Trimis JL. Associations of electronic cigarette nicotine concentration with subsequent cigarette smoking and vaping levels in adolescents. JAMA Pediatr. 2017; 171(12): 1192–9.

11. Jackson CB, Farzan M, Chen B, Choe H. Mechanisms of SARS-CoV-2 entry into cells. Nature Reviews Molecular Cell Biology. 2022;23(1): 3–20.

12. Hoffmann M, Kleine-Weber H, Schroeder S, Krüger N, Herrler T, Erichsen S, Schiergens TS, Herrler G, Wu N-H, Nitsche A, Müller M A, Drosten C, Pöhlmann S. SARS-CoV-2 cell entry depends on ACE2 and TMPRSS2 and is blocked by a clinically proven protease inhibitor. Cell. 2020;181(2):271-280.e8.

13. Gaiha SM, Cheng J, Halpern-Felsher B. Association between youth smoking, electronic cigarette use, and COVID-19. Journal of Adolescent Health. 2020 Oct 1;67(4):519–23.

14. Li D, Croft DP, Ossip DJ, Xie Z. The association between statewide vaping prevalence and COVID-19. Preventive Medicine Reports. 2020;20:101254

15. Wang Q, Sundar IK, Li D, Lucas JH, Muthumalage T, McDonough SR, et al. E-cigarette-induced pulmonary inflammation and dysregulated repair are mediated by nAChR α7 receptor: role of nAChR α7 in SARS-CoV-2 Covid-19 ACE2 receptor regulation. Respir Res. 2020;21(1):154.

16. Naidu V, Zeki AA, Sharma P. Sex differences in the induction of angiotensin converting enzyme 2 (ACE-2) in mouse lungs after e-cigarette vapor exposure and its relevance to COVID-19. J Investig Med. 2021;69(5):954–61

17. Masso-Silva JA, Moshensky A, Shin J, Olay J, Nilaad S, Advani I, et al. Chronic E-cigarette aerosol inhalation alters the immune state of the lungs and increases ACE2 expression, raising concern for altered response and susceptibility to SARS-CoV-2. Front Physiol. 2021;12:649604.

18. Lallai V, Manca L, Fowler CD. E-cigarette vape and lung ACE2 expression: Implications for coronavirus vulnerability. Environmental Toxicology and Pharmacology. 2021;86:103656.

19. Ghosh A, Coakley RC, Mascenik T, Rowell TR, Davis ES, Rogers K, et al. Chronic e-cigarette exposure alters the human bronchial epithelial proteome. Am J Respir Crit Care Med. 2018;198(1):67–76.

20. McAlinden KD, Lu W, Ferdowsi PV, Myers S, Markos J, Larby J, et al. Electronic cigarette aerosol is cytotoxic and increases ACE2 expression on human airway epithelial cells: implications for SARS-CoV-2 (COVID-19). JCM 2020;10(5):1028.

21. Phandthong R, Wong M, Song A, Martinez T, Talbot P. New insights into how JUUL™ electronic cigarette aerosols and aerosol constituents affect SARS-CoV-2 infection of human bronchial epithelial cells. bioRxiv 2022.08.23.505031 [preprint]. 2022 August 24. Available from https://www.biorxiv.org/content/10.1101/2022.08.23.505031v1

22. Jose T, Croghan IT, Hays JT, Schroeder DR, Warner DO. Electronic cigarette use is not associated with COVID-19 diagnosis. J Prim Care Community Health 2021;12:215013272110243.

23. Burnett-Hartman AN, Goldberg Scott S, Powers JD, Clennin MN, Lyons JA, Gray M, et al. The association of electronic cigarette use with SARS-CoV-2 infection and COVID-19 disease severity. Tob Use Insights. 2022;15:1179173×2210966.

24. Shrimp JH, Kales SC, Sanderson PE, Simeonov A, Shen M, Hall MD. An enzymatic TMPRSS2 assay for assessment of clinical candidates and discovery of inhibitors as potential treatment of COVID-19. ACS Pharmacology TranslationalScience. 2020;3(5):997–1007.

25. Crawford KHD, Eguia R, Dingens AS, Loes AN, Malone KD, Wolf CR, Chu HY, Tortorici, MA, Veesler D, Murphy M, Pettie D, King NP, Balazs AB, Bloom JD. protocol and reagents for pseudotyping lentiviral particles with SARS-CoV-2 spike protein for neutralization assays. Viruses. 2020;12(5): 513.

26. Behar RZ, Hua M, Talbot P. Puffing topography and nicotine intake of electronic cigarette users. PLOS ONE. 2015;10(2):e0117222.

27. Pankow JF, Kim K, McWhirter KJ, Luo W, Escobedo JO, Strongin RM, Duell AK, Peyton DH. Benzene formation in electronic cigarettes. PLOS ONE 2017;12(3), e0173055.

28. Omaiye EE, McWhirter KJ, Luo W, Pankow JF, Talbot P. High-Nicotine Electronic Cigarette Products: Toxicity of JUUL Fluids and Aerosols Correlates Strongly with Nicotine and Some Flavor Chemical Concentrations. Chem Res Toxicol. 2019;32(6):1058–1069.

29. Talih S, Salman R, El-Hage R, Karam E, Karaoghlanian N, El-Hellani A, Saliba N, Shihadeh A. (2019). Characteristics and toxicant emissions of JUUL electronic cigarettes. Tobacco Control. 2019;28(6):678–80.

30. Williams M, Li J, Talbot P. Effects of model, method of collection, and topography on chemical elements and metals in the aerosol of tank-style electronic cigarettes. Scientific Reports. 2019;9(1):13969.

31. Son Y, Mishin V, Laskin JD, Mainelis G, Wackowski OA, Delnevo C, Schwander S, Khlystov A, Samburova V, Meng Q. Hydroxyl radicals in e-cigarette vapor and e-vapor oxidative potentials under different vaping patterns. Chemical Research in Toxicology. 2019;32(6):1087–95.

32. Zhao D, Navas-Acien A, Ilievski V, Slavkovich V, Olmedo P, Adria-Mora B, Domingo-Relloso A, Aherrera A, Kleiman NJ, Rule AM, Hilpert M. Metal concentrations in electronic cigarette aerosol: Effect of open-system and closed-system devices and power settings. Environmental Research. 2019;174:125–34.

33. Dautzenberg, B. Real-time characterization of e-cigarettes use: the 1 million puffs study. Journal of Addiction Research & Therapy. 2015;06(02).

34. Bitzer, ZT, Goel R, Reilly SM, Foulds J, Muscat J, Elias RJ, Richie JP Effects of solvent and temperature on free radical formation in electronic cigarette aerosols. Chemical Research in Toxicology. 2018;31(1): 4–12.

35. Erythropel HC, Davis LM, de Winter TM, Jordt SE, Anastas PT, O’Malley SS, Krishnan-Sarin S, Zimmerman, JB. Flavorant–solvent reaction products and menthol in juul e-cigarettes and aerosol. American Journal of Preventive Medicine. 2019;57(3):425–27.

36. Jensen RP, Strongin RM, Peyton DH. Solvent chemistry in the electronic cigarette reaction vessel. Scientific Reports. 2017;7(1):42549.

37. Williams M, Bozhilov K, Ghai S, Talbot P. Elements including metals in the atomizer and aerosol of disposable electronic cigarettes and electronic hookahs. PLOS ONE. 2017;12(4): e0175430.

38. PubChem [Internet]. Bethesda (MD): National Library of Medicine (US), National Center for Biotechnology Information; 2004-. PubChem Compound Summary for CID 243, Benzoic acid; [accessed 2022 July 29]. Available from: https://pubchem.ncbi.nlm.nih.gov/compound/Benzoic-acid.

39. Cunningham A, McAdam K, Thissen J, Digard H. The evolving e-cigarette: comparative chemical analyses of e-cigarette vapor and cigarette smoke. Front Toxicol. 2020;2:586674.

40. Scholtissek, C., Bürger, H., Kistner, O., & Shortridge, K. F. (1985). The nucleoprotein as a possible major factor in determining host specificity of influenza H3N2 viruses. Virology. 147(2), 287–294.

41. Zhou T, Tsybovsky Y, Gorman J, Rapp M, Cerutti G, Chuang GY, Katsamba PS, Sampson JM, Schön A, Bimela J, Boyington JC, Nazzari A, Olia AS, Shi W, Sastry M, Stephens T, Stuckey J, Teng IT, Wang P, Wang S, Zhang B, Friesner RA, Ho DD, Mascola JR, Shapiro L, Kwong PD. Cryo-EM structures of SARS-CoV-2 spike without and with ACE2 reveal a pH-dependent switch to mediate endosomal positioning of receptor-binding domains. Cell Host Microbe. 2020;28(6):867-879.e5.

42. Xie Y, Guo W, Lopez-Hernadez A, Teng S, Li L. The pH effects on SARS-CoV and SARS-CoV-2 spike proteins in the process of binding to hACE2. Pathogens. 2022;11(2): 238.

43. Luo B, Schaub A, Glas I, Klein LK, David SC, Bluvshtein N, Violaki K, Motos G, Pohl M, Hugentobler W, Nenes A, Krieger UK, Stertz S, Peter T, Kohn T. Acidity of expiratory aerosols controls the infectivity of airborne influenza virus and SARS-CoV-2. medRxiv 2022.03.14.22272134 [preprint]. 2022 June 21. Available from https://www.medrxiv.org/content/10.1101/2022.03.14.22272134v2

44. Stefaniu A, Pirvu L, Albu B, Pintilie L. Molecular docking study on several benzoic acid derivatives against SARS-CoV-2. Molecules. 2020;25(24):5828.

45. Rocchitta G, Bacciu A, Arrigo P, Migheli R, Bazzu G, Serra PA. Propylene glycol stabilizes the linear response of glutamate biosensor: potential implications for in-vivo neurochemical monitoring. Chemosensors. 2018;6(4):58.

46. Zhang L, Jackson CB, Mou H, Ojha A, Peng H, Quinlan BD, Rangarajan ES, Pan A, Vanderheiden A, Suthar MS, Li W, Izard T, Rader C, Farzan M, Choe H. SARS-CoV-2 spike-protein D614G mutation increases virion spike density and infectivity. Nature Communications. 2020;11(1): 6013

47. Woodall M, Jacob J, Kalsi KK, Schroeder V, Davis E, Kenyon B, et al. E-cigarette constituents propylene glycol and vegetable glycerin decrease glucose uptake and its metabolism in airway epithelial cells in vitro. American Journal of Physiology-Lung Cellular and Molecular Physiology. 2020;319(6):L957–67.

48. National Academies of Sciences E, and Medicine; Health and Medicine Division; Board on Population Health and Public Health Practice; Committee on the Review of the Health Effects of Electronic Nicotine Delivery Systems. Public Health Consequences of E-Cigarettes. In: KL Edl, Stratton K, editors. Toxicology of E-Cigarette Constituents. Volume 5, edn. Washington (DC): National Academies Press (US); 2018.

49. Gheware A, Ray A, Rana D, Bajpai P, Nambirajan A, Arulselvi S, Mathur P, Trikha A, Arava S, Das P, Mridha AR, Singh G, Soneja M, Nischal N, Lalwani S, Wig N, Sarkar C, Jain D. ACE2 protein expression in lung tissues of severe COVID-19 infection. Scientific Reports. 2022;12(1): 4058.

50. Patanavanich R, Glantz SA. Smoking is associated with COVID-19 progression: a meta-analysis. Nicotine & Tobacco Research. 2020;22(9):1653–56.

51. Muus C, Luecken MD, Eraslan G, Sikkema L, Waghray A, Heimberg G, Kobayashi Y, Vaishnav ED, Subramanian A, Smillie C, Jagadeesh KA, Duong ET, Fiskin E, Torlai Triglia E, Ansari M, Cai P, Lin B, Buchanan J, Chen S, et al. Single-cell meta-analysis of SARS-CoV-2 entry genes across tissues and demographics. Nature Medicine. 2021;27(3): 546–59.

52. Dorjee, K., Kim, H., Bonomo, E., & Dolma, R. (2020). Prevalence and predictors of death and severe disease in patients hospitalized due to COVID-19: A comprehensive systematic review and meta-analysis of 77 studies and 38,000 patients. PLOS ONE. 2020;15(12):e0243191.

53. Changeux JP, Amoura Z, Rey FA, Miyara M. A nicotinic hypothesis for Covid-19 with preventive and therapeutic implications. C R Biol. 2020;343(1):33–39

54. Farsalinos K, Bagos PG, Giannouchos T, Niaura R, Barbouni A, Poulas K. Smoking prevalence among hospitalized COVID-19 patients and its association with disease severity and mortality: An expanded re-analysis of a recent publication. Harm Reduct J. 2021;18(1):9.

## Supplement file references

1. Pankow JF, Kim K, McWhirter KJ, Luo W, Escobedo JO, Strongin RM, Duell AK, Peyton DH. Benzene formation in electronic cigarettes. PLOS ONE 2017;12(3), e0173055.

2. Omaiye EE, McWhirter KJ, Luo W, Pankow JF, Talbot P. High-Nicotine Electronic Cigarette Products: Toxicity of JUUL Fluids and Aerosols Correlates Strongly with Nicotine and Some Flavor Chemical Concentrations. Chem Res Toxicol. 2019;32(6):1058–1069

3. Crawford KHD, Eguia R, Dingens AS, Loes AN, Malone KD, Wolf CR, Chu HY, Tortorici, MA, Veesler D, Murphy M, Pettie D, King NP, Balazs AB, Bloom JD. protocol and reagents for pseudotyping lentiviral particles with SARS-CoV-2 spike protein for neutralization assays. Viruses. 2020;12(5): 513.

